# Swashing motility: A novel propulsion-independent mechanism for surface migration in *Salmonella* and *E. coli*

**DOI:** 10.1101/2024.08.21.609010

**Authors:** Justin Panich, Eric M. Dudebout, Navish Wadhwa, David F. Blair

## Abstract

Bacterial motility over surfaces is crucial for colonization, biofilm formation, and pathogenicity. Surface motility in *Escherichia coli* and *Salmonella enterica* is traditionally believed to rely on flagellar propulsion. Here, we report a novel mode of motility, termed “swashing,” where these bacteria migrate on agar surfaces without functional flagella. Mutants lacking flagellar filaments and motility proteins exhibit rapid surface migration comparable to wild-type strains. Unlike previously described sliding motility, swashing is inhibited by surfactants and requires fermentable sugars. We propose that the fermentation of sugars at the colony edge produces osmolytes, creating local osmotic gradients that draw water from the agar, forming a fluid bulge that propels the colony forward. Our findings challenge the established view that flagellar propulsion is required for surface motility in *E. coli* and *Salmonella*, and highlight the role of a fermentation in facilitating bacterial spreading. This discovery expands our understanding of bacterial motility, offering new insights into bacterial adaptive strategies in diverse environments.

**Significance Statement:** Bacteria move on surfaces using a variety of mechanisms, with important implications for their growth and survival in both the clinical setting (such as on the surface of medical devices) and in the wild. Surface motility in the medically important model species *S. enterica* and *E. coli* has been extensively studied and is thought to require flagellar propulsion. Here, we show surface expansion in these species even in the absence of propulsion by the flagella. Instead, movement is tied to fermentation and surface tension: As cells ferment sugars, they create local osmolarity gradients, which generate a wave of fluid on which the cells “swash.”

## Introduction

The ability to move allows many species of bacteria to seek and colonize favorable environments, avoid unfavorable conditions, and form complex macroscopic structures such as biofilms or fruiting bodies (*1–4*). Mechanisms of bacterial motility have been intensively studied and can be grouped into different classes based on underlying mechanisms and the setting (e.g., liquid vs solid media) where the behavior occurs (*1, 5*). Many species swim in liquid media using flagella (Fig. 1*A*), which harness the membrane ion gradient to drive rotation of helical filaments that function as propellers (*6–9*). Movement on surfaces occurs by diverse mechanisms, including the extension and retraction of specialized pili (“twitching”) (*10*), the movement of membrane-bound adhesins along the cell surface (“gliding”) (*11*), by surfactant-assisted but largely passive growth-based mechanisms (“sliding”) (*12*), or by swimming within a thin layer of fluid on the surface (“swarming”) (*13–15*). Sliding and swarming on agar surfaces have been observed in diverse genera, including *Proteus* (*16*), *Vibrio* (*17*), *Bacillu*s (*18*), and Clostridi*um* (*19*), as well as in the gram-negative model species *Escherichia coli* and *Salmonella typhimurium* (*15*).

**Fig. 1.**
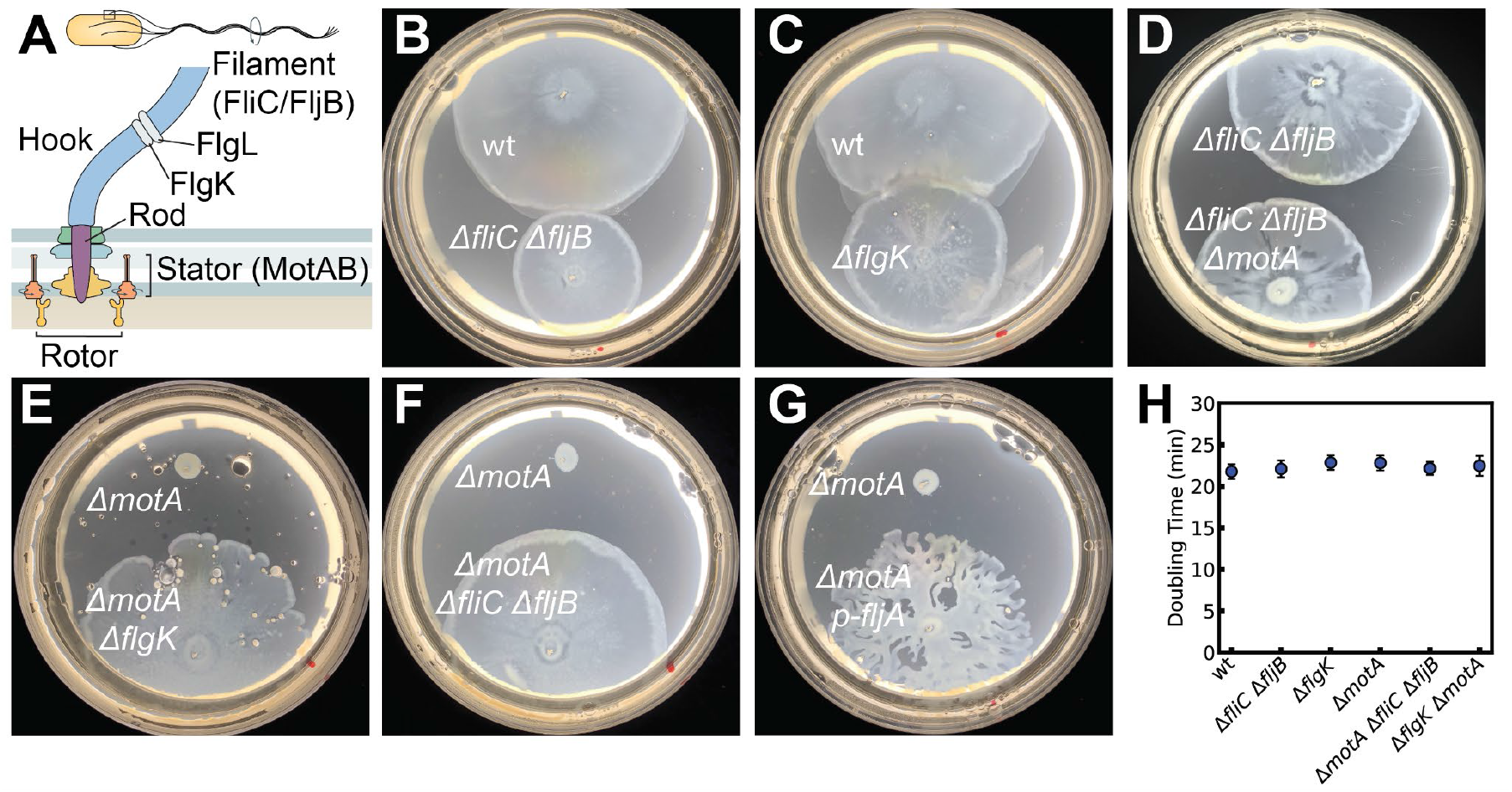
Propulsion-independent surface migration in *S. enterica LT2*. (*A*) Schematic of the bacterial flagellar motor. Stators (MotAB) power the rotation of the rotor, which is connected to a flexible hook by a rigid rod. This hook is attached to the flagellar filament (FliC/FljB) through the hook-associated proteins FlgK and FlgL. (*B-D*) Migration of both motile (w.t.) and immotile strains on swarm plates (LB, 0.5% glucose, 0.55% Bacto-agar). Plates were spotted with 4 µl of overnight culture and incubated at 37° C for 10 h (B and C) or 12 h (*D*). (*E-G*) Failure of an immotile Δ*motA* strain to migrate, and rescue of migration by removal of the flagellar filament either by deletion of the hook-filament junction protein (Δ*flgK*), deletion of the flagellin genes (Δ*fliC*Δ*fljB*), or inhibition of flagellin translation by overexpression of the regulator FljA. Plates were spotted with 4 µl of overnight culture and incubated at 37° C for 12 h. (*H*) Doubling time of the *Salmonella* strains when grown in LB + 0.5% (w/v) at 37° C, from 100-fold dilutions of an overnight culture.

While swarming by *S. enterica* and *E. coli* has been studied extensively (*20–30*), its mechanism is not completely understood. *S. enterica* and *E. coli* are termed “temperate” swarmers because their movement requires relatively low agar concentrations (∼0.55%), in contrast to the robust swarmer *Proteus mirabilis*, which can migrate on relatively firm (>2% agar) surfaces (*15, 31*). The agar formulation is also important; *E. coli* swarms much better on Eiken-brand agar than on other brands (*15*). Whereas swarming of *Bacillus subtilis, Serratia marcescens*, and *Pseudomonas aeruginosa* depends on cell-produced surfactants (*13, 29, 32–34*), movement of *E. coli* and *S. enterica* does not involve surfactant but relies on cell-derived osmolytes that draw water out of the plate to form a wet zone at the margin of the expanding colony (*27, 29, 35, 36*). The relevant osmolyte(s) has not been identified. Movement of the cells within this fluid layer is thought to depend on active propulsion by the flagella; *S. enterica* cells deleted for the flagellin proteins that form the filament, or the motility proteins MotA and MotB needed for flagella rotation, reportedly fail to migrate (*15*).

Here, we show that propulsion by flagella is not required for surface migration of *Salmonella* and *E. coli*. Though immotile in liquid, mutants lacking the flagellar filament moved on swarm plates at more than half the rate of wild-type cells. Cells lacking filament and the motility protein MotA also migrated at substantial rates. The movement described here is different from the movement reported previously for certain immotile *Salmonella* and *E. coli* strains on agarose (*37, 38*); it is more rapid, occurs on standard swarm plates, and does not require the protein PagM. It is also different from the sliding motility seen in other species, as it does not involve cell-produced surfactant and is instead inhibited by adding a surfactant (Tween-80) at concentrations that accelerate the swarming of motile, wild-type cells (*39*). Movement requires the presence of fermentable sugar, and migrating cells produce acid and fermentation products in their wake. We rationalize these results using a model in which the production of osmolytes by fermentation and resulting water movements generate a moving bulge of fluid at the margin of the expanding colony. Because it involves cells riding a wave of fluid, this mode of surface movement can be called “swashing.”

## Results

### Propulsion-independent surface movement in *Salmonella*

We discovered swashing motility in an experiment first intended as a negative control. In this experiment, we examined the migration of a Salmonella strain, which had both flagellin genes *fliC* and *fljB* deleted, on swarm plates (LB, 0.5% glucose, 0.55% agar). The Δ*fliC*Δ*fljB* mutant assembles the hook-basal body (HBB) but lacks the flagellar filament required for propulsion (Fig. 1*A*). Surprisingly, the immotile Δ*fliC*Δ*fljB* cells migrated on swarm plates at more than half the rate of motile wild-type cells (Fig. 1*B*). Cells grown in liquid culture were immotile when examined under the microscope (Fig. S1) and, as expected, failed to migrate in softer (0.3% agar) motility plates (Fig. S2).

Prompted by this finding, we examined other non-swimming mutants. A Δ*flgK* mutant, which lacks the hook-filament junction and assembles only the HBB, also migrated rapidly on swarm plates (Fig. 1*C*). Like the Δ*fliC*Δ*fljB* mutant, the Δ*flgK* mutant was immotile when examined under the microscope (Fig. S1) and failed to migrate in 0.3%-agar motility plates (Fig. S2). A Δ*flgL* mutant, which also only assembles the HBB, displayed the same phenotype as Δ*flgK* (Fig. S3). Thus, mutants that assemble just the HBB, though immotile in liquid and soft-agar “swim” plates, migrate fairly rapidly on the surface of swarm plates. The hook structure still present in these mutants might, in principle, enable some propulsion. To test this possibility, we prevented hook rotation by deleting the motility gene

*motA*, in the filamentless Δ*fliC*Δ*fljB* background. Migration was not impaired by paralysis of the hook (Fig. 1*D*).

The rapid migration of the Δ*motA* strain lacking the filament contrasts with the non-migrating phenotype of a Δ*motA* mutant reported previously by Harshey and coworkers (*15*) and reproduced here (Fig. 1 *E-G*). Comparisons in Fig. 1 *F-H* show that migration of the Δ*motA* mutant is rescued when assembly of the filament is prevented, either by deletion of the flagellin genes, deletion of *flgK*, or overexpression of a regulatory protein (FljA) that inhibits translation of *fliC*. Thus, the failure of the Δ*motA* strain to migrate appears to be due to the presence of paralyzed filaments, rather than the absence of active propulsion. The differences in expansion rates of these various strains were not correlated to any differences in their growth rates (Fig. 1*H*).

### Propulsion-independent surface movement in *E. coli*

*E. coli* also swarms on agar plates, most robustly with a particular (Eiken-brand) agar formulation (*15*). To examine the generality of the propulsion-independent surface migration, we repeated the experiments described above for *Salmonella* with *E. coli* K12 (strain MG1665). With nutrient sources like those used for *Salmonella* (LB supplemented with 0.5% glucose), w.t. *E. coli* swarmed reproducibly at rates similar to w.t. *Salmonella* (Fig. 2*A*). As in *Salmonella, E. coli* mutants lacking the flagellar filament, whether due to loss of flagellin FliC or the hook-filament connector FlgK, retained the ability to move at a significant rate on swarm plates (Fig. 2 *B,C*). Δ*flgK* and Δ*fliC* mutant cells grown in liquid culture were immotile when examined by light microscopy (Fig. S4), and did not migrate in soft-agar (0.30%) swim plates (Fig. S5). Thus, like *S. enterica, E. coli* migrates on swarm plates in a process that does not require propulsion by the flagella.

**Fig. 2.**
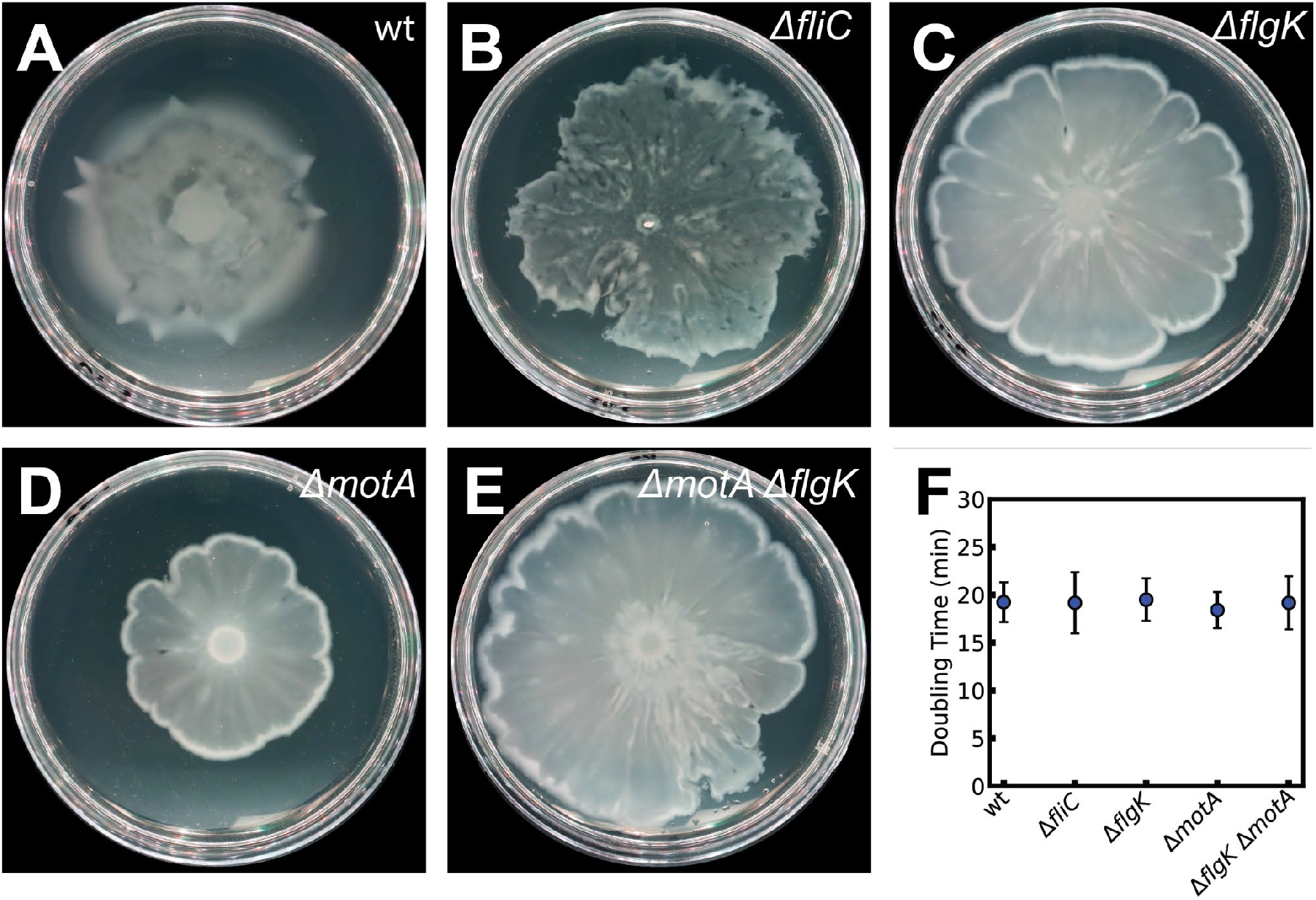
Propulsion-independent surface migration in *E. coli*. (*A-C*) Migration of the wild type, and filamentless mutants defective in either flagellin (FliC) or the hook-junction filament protein FlgK. Plates contained LB, 0.5% glucose, and 0.5% Eiken agar. (*D and E*) Reduction in migration upon deletion of *motA*, and rescue upon further deletion of *flgK*. After spotting with 2 µl of overnight culture, plates were incubated at 37° C for 12 h (*A*) and 16 h (*B-E*). (*F*) Doubling time of the *E. coli* strains when grown in LB + 0.5% (w/v) at 37° C, from 100-fold dilutions of an overnight culture.

Next, we examined swashing in *E. coli* mutants that lacked the motility protein MotA. Migration significantly decreased, but did not entirely cease, upon deletion of *motA* (Fig. 2*D*). As in *Salmonella*, rapid migration in the Δ*motA* mutant was rescued by further deletion of *flgK* (Fig. 2E). Both the Δ*motA* and Δ*motAΔflgK* strains were immotile in liquid (Fig. S4) and in soft-agar swim plates (Fig. S5). The differences in expansion rates of the various strains did not correlate with any differences in their growth rates (Fig. 2*F*).

### Surfactants inhibit swashing in both *Salmonella* and *E. coli*

Propulsion-independent migration on surfaces has been seen in other species previously, such as *Pseudomonas* (*40*), and in those cases, it has been termed sliding (*12*). In documented cases of sliding motility, the movement has been shown to involve surfactants or certain cell-surface molecules that facilitate movement (*38*). The propulsion-independent movement described here does not involve surfactant: Migration of the Δ*flgK* strain persisted in a *Salmonella* Δ*srfB* mutant defective in surfactant production (Fig. S6), and in both *Salmonella* and *E. coli*, swashing was inhibited rather than helped by the surfactant Tween-80 (Fig. 3 and Fig. S7). Swarming of the wild-type strains of both species was, by contrast, enhanced by Tween-80 (Fig. 3 and Fig. S7).

**Fig. 3.**
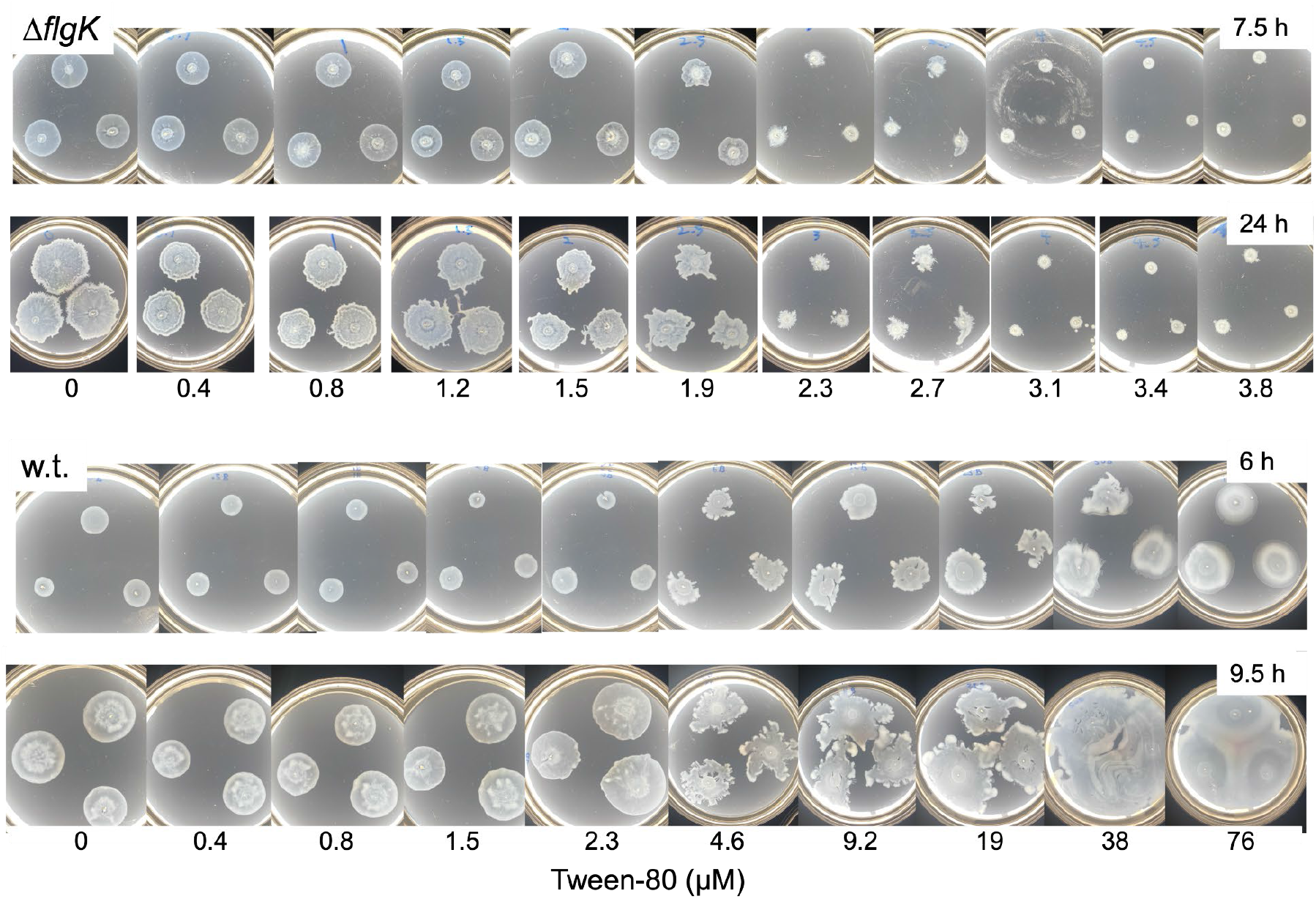
Inhibition of *Salmonella* swashing and enhancement of swarming by surfactant. Tween-80 was added to the concentrations indicated, during pouring. Plates were spotted with 4 µl of overnight culture and incubated at 37° C for the indicated times.

Tween-80 did not greatly affect the swashing of Salmonella Δ*flgK* at levels below 1.5 mM. At 3 mM and above, it prevented swashing. At concentrations of about 2.5 mM, colonies displayed irregular shapes, suggesting the occurrence of abortive swashing at certain locations (Fig. 3). Migration of w.t. cells (i.e., swarming) was not prevented by Tween-80 at any concentration tested (Fig. 3). Irregular colony shapes were seen at intermediate concentrations (around 10 mM), and higher concentrations caused faster swarming. These trends were similar in *E. coli* (Fig. S7). Below 1 µM Tween-80, *E. coli* swashing was unaffected. Above this concentration, *E. coli* swashed with irregular morphology and dose-dependent reduction in final colony diameters. In contrast, and in agreement with prior work (*39*), swarming in w.t. *E. coli* was enhanced by increasing Tween-80 concentrations (Fig. S7). Tween-80 did not affect cell growth at concentrations tested here, suggesting that the inhibition of swashing by Tween-80 was not caused by any growth defects (Fig. S8).

### Swashing is associated with fermentation

For reasons that have not been clear, glucose has been found essential for swarming in *Salmonella* and *E. coli* and is typically included in plates at a concentration of 0.5% (*15, 41*). Consistent with previous reports (*15*), we observed that swarming of wild-type *Salmonella* and *E. coli* ceased at glucose concentrations below 0.25% and 0.3%, respectively. Migration of the Δ*flgK* strain in both species showed a similar dependence on glucose (Fig. S9, Fig. S10). This prompted us to investigate the effect of various sugars on swashing. At the standard concentration of 0.5%, maltose and xylose supported swashing of the *Salmonella ΔflgK* strain at significant rates (Fig. 4*A*). Arabinose and galactose, and to a smaller extent sucrose, gave rise to small protrusions. When plates were supplemented with 0.2% glucose (a concentration insufficient to enable migration by itself), swashing was strongly enhanced in plates with maltose, xylose, arabinose, or galactose (Fig. 4*A*).

**Fig. 4.**
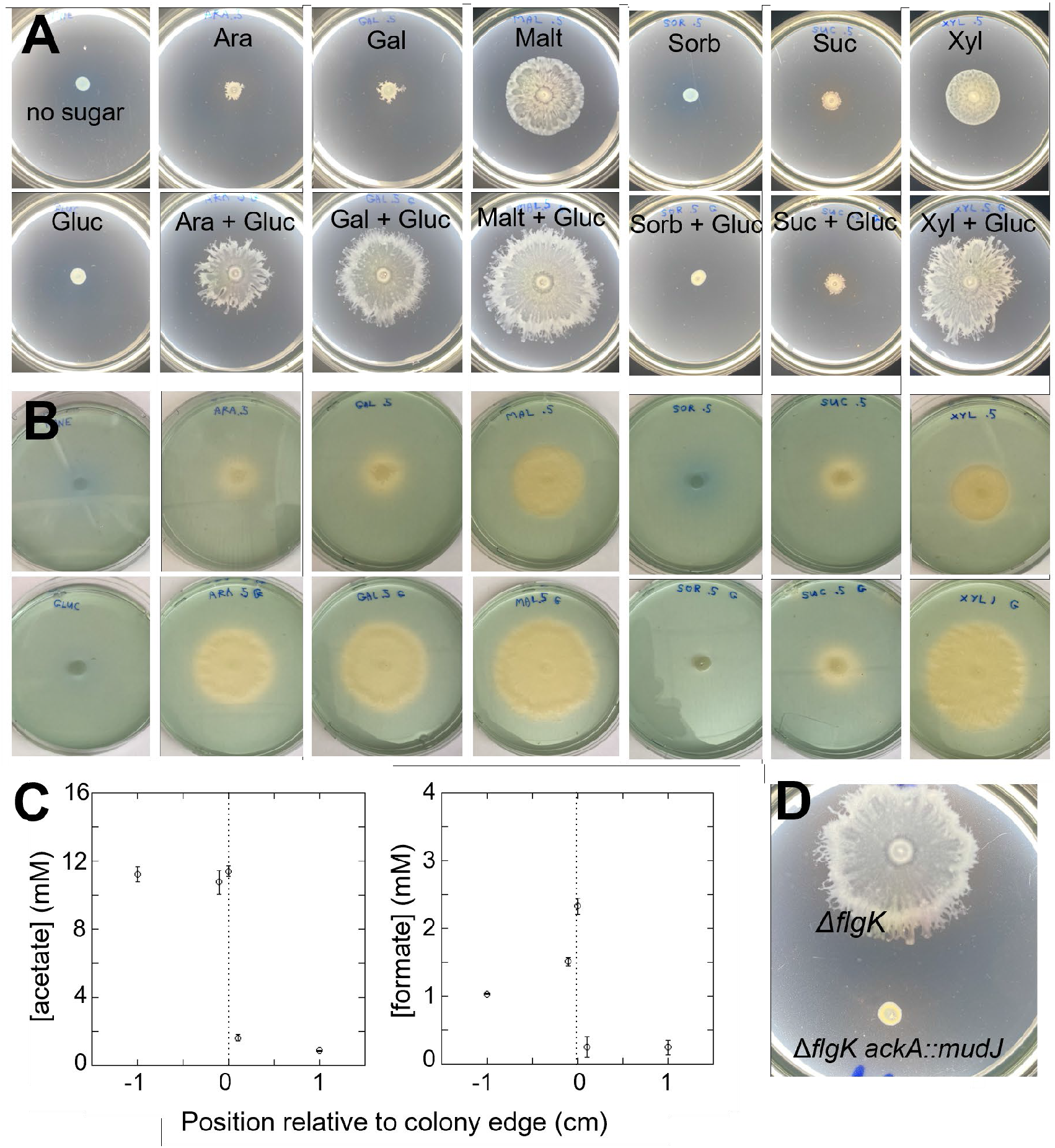
Association of swashing with fermentation. (*A*) Effects of various sugars on migration of the Δ*flgK* strain, and (*B*) acidification of the plates by the advancing cells. The Δ*flgK* strain was spotted on swarm plates containing the indicated sugar sources and the pH indicator bromothymol blue (1.25 % (v/v) of 1 mg/mL). Where indicated, glucose was added to a concentration of 0.2%; other sugars were present at the standard concentration of 0.5%. Plates were spotted with 4 µl of overnight culture and incubated at 37° C for 16.5 h. In (*B*), the same plates were imaged to highlight the color change due to acidification. (*C*) Production of acetate and formate by the advancing cells. (*D*) Effect of deleting *ackA* (encoding acetate kinase) on migration of the Δ*flgK* strain. The plate was incubated at 37° C for 17 h.

The sugars that support swashing motility are all fermentable by *Salmonella*, and at the high cell densities occurring on plates, some fermentation might be expected to occur (owing to sub-optimal oxygenation) and should acidify the plates. We observed acidification in plates containing the indicator bromothymol blue (1.25 % (v/v) of 1 mg/mL) for any of the fermentable sugars (Fig. 4*A*). The mixed-acid fermentation carried out by *Salmonella* should produce acetate and formate as major products (*42, 43*). Both products were seen in the wake of migrating cells (Fig. 4*B*). While the glucose in the plate was only partially utilized by cells at the colony margin, glucose concentration was decreased across the margin by about half the increase in acetate, as expected if fermentation is the major pathway for glucose utilization. Acetate concentration was elevated and fairly uniform behind the advancing front, whereas formate spiked near the colony margin, consistent with the reuptake and subsequent oxidation of formate that has been documented in *E. coli* and *Salmonella* (*44, 45*). Disruption of the gene for acetate kinase, required for mixed-acid fermentation, prevented migration of the Δ*flgK* strain (Fig 4*C*).

## Discussion

The ability of *E. coli* and *Salmonella* to migrate rapidly on surfaces without flagellar propulsion challenges previous findings. Swarming was reportedly prevented by deleting the two flagellin genes in *Salmonella*, and mutants with defects in the motility genes *motA* or *motB* also failed to swarm (*15*). Furthermore, even in cells with rotating flagella, swarming requires CW/CCW reversals of the flagellar motor (*28*). Microarray analysis revealed the upregulation of late flagellar promoters in swarming cells (*20*), suggesting flagellar propulsion is essential for swarming. Our study demonstrates, however, that flagellar propulsion is dispensable for the migration of *E. coli* and *Salmonella* on soft agar surfaces.

Whereas the rapid movement of filament-less strains (Fig. 1) contrasts with earlier reports, the reduced mobility of Δ*motA* cells is consistent with previous findings (*15*). Notably, the outward migration of the Δ*motA* strain can be restored if the filament is also removed, suggesting that non-rotating filaments impede migration. Paralyzed filaments might interact with the surface or nearby cells to inhibit movement. Friction due to nonrotating filaments does not appear to hinder cell movement strongly, however, because Δ*motA* cells moved readily on the surface when present in mixed cultures alongside robustly swashing Δ*flgK* cells (Fig. S11).

We also considered the effects of filament loss on non-reversing chemotactic knockout strains. Consistent with previous reports (*46*), Δ*cheY Salmonella* cells, locked in CCW rotation, fail to migrate, but surface motility was restored by further deletion of the filament (Fig. S12). Our observations suggest that some aspects of flagellar function remain important for migration, even though propulsion is dispensable. We speculate that the basal body and hook, which form a channel that carries various protein cargoes during flagellar export, might conduct osmolytes out of the cell. Mutations that remove the filament may allow increased osmolyte secretion under certain circumstances (such as when the filament is paralyzed).

Swashing is distinct from other forms of propulsion-independent mechanisms of motility described in the literature. A previous report described propulsion-independent surface migration in *Salmonella* on 0.3% agarose plates, dependent on the protein PagM (*38*). The migration described here is different: it occurs on 0.55% agar, is faster, and does not require PagM, which is absent in the *Salmonella* strains (LT2 and derivatives) studied here. Swashing is also distinct from the “sliding” motility observed in several species (*12*). Sliding, driven by cell growth, is facilitated by surfactants that are not produced by the strains studied here. Moreover, the swashing motility we observed is inhibited by adding surfactant, contrasting with previously reported sliding in *Salmonella* and *E. coli* (*37, 38*). In a colony that is expanding by a growth-based mechanism, the density of cells should remain high behind (i.e., inside) the advancing edge of the colony. This contrasts with what is seen in expanding colonies of the (swashing) Δ*flgK* strain, where a thin zone of high cell density advances outward while leaving a zone of lower density behind (Fig. S13). In a model proposed below, swashing motility depends on a fluid bulge at the margin of the colony; we suggest that the decreased surface tension resulting from surfactant addition hinders the formation of this bulge.

The requirement for a fermentable sugar, the acidification of plates behind swashing cells, and the accumulation of fermentation by-products all point to an association between swashing motility and fermentation. The acetate and formate concentrations shown in Fig. 4*B* likely underestimate the actual concentrations in the fluid layer containing the cells, as they were measured using plugs that sampled the entire thickness of the agar. Moreover, actual formate concentrations at the colony margin are likely to be higher than measured, as the spiked shape of the formate profile (Fig. 4B) suggests the re-acquisition of secreted formate. Nevertheless, the measured total acetate and formate (about 15 mM) represents a significant fraction of the osmolarity of LB-glucose swarm medium (around 250 mM) (*47*). Thus, the secreted acetate and formate should significantly affect local osmolarity. We suggest that acetate and formate are the osmolytes implicated (but not previously identified) in the surface and swarming motility of *Salmonella* and *E. coli* (*15*).

Figure 5 presents a model for swashing motility. Movement is associated with a bulge of fluid and increased cell density at the margin of the expanding colony. This bulge is generated by elevated osmolarity due mainly to the production of acetate and formate by cells near the outer edge. The influx of fluid pushes the colony front forward. Behind the bulge, osmolyte production is slower and the osmolytes diffuse back into the agar. This model explains the role of glucose in the media used for surface migration studies of *Salmonella* and *E. coli*. Besides supporting growth, glucose fermentation also drives osmotic flows that hydrate the colony front, facilitating expansion. Glucose has been shown to also enhance propulsion-independent surface movement in Mycobacteria (*48*), suggesting that a similar mechanism might be at play there.

**Fig. 5.**
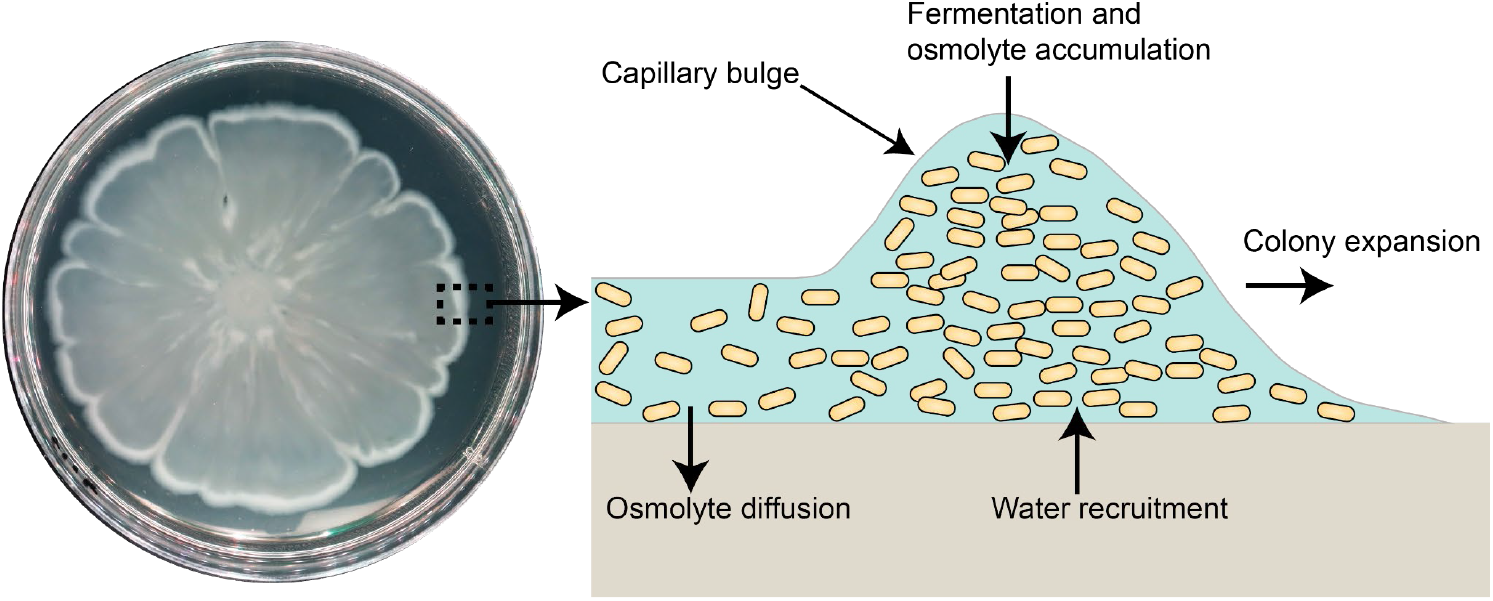
Model for bacterial swashing, showing a cross-section of the colony edge. Bacterial fermentation of sugars generates osmolytes, creating an osmotic gradient at the colony’s leading edge. This imbalance draws water from the agar, forming a capillary bulge and driving outward expansion. Behind the front, both osmolytes and water diffuse into the agar. Cells are carried forward by the resulting solvent bulge at the colony margin.

This model is similar to the one proposed by Wu and Berg (*27*), with a key distinction. While Wu and Berg suggested that flagellar rotation pumps fluid outward at the swarm front, we propose that colony expansion is primarily driven by osmotically induced fluid influx. Bru *et al*. (*49*) recently questioned the role of flagellar propulsion in *P. aeruginosa* swarm expansion, offering arguments that align with our findings. Our model also parallels that of Srinivasan *et al*. for *Bacillus subtilis* colonies, where expansion is mainly driven by osmotic flux without explicit contribution from flagellar motility (*50*). However, unlike their findings of a power-law dependence on surface tension, we observed a sharp transition from expansion to no expansion with the concentration of Tween-80 (and thus surface tension) in the medium (Fig. 3).

In summary, we describe a novel mode of bacterial translocation that does not rely on active propulsion but instead utilizes osmotic flows generated by the products of fermentative metabolism. Since key components of this motility—fermentation and semi-permeable wet surfaces—are widespread, we hypothesize that this mechanism significantly contributes to the range expansion of many bacterial species and other single-celled organisms.

## Materials and Methods

### Bacterial strains and cultures

The *Salmonella* and *E. coli* strains used in this study are listed in Tables S1 and S2, respectively. *E. coli* and *Salmonella* strains were grown in LB liquid media at 37 °C with appropriate antibiotics. When plating on solid media, LB media was solidified with 1.5 % (w/v) agar, supplemented with appropriate antibiotics, at concentrations of 10 µg/mL for tetracycline, 100 µg/mL for ampicillin, and 50 µg/mL for spectinomycin.

### Strains and Vector Construction

Karlinsey’s Lambda red recombineering method (*51*) was used to create the knockout strains of *E. coli* and *Salmonella* used here. Briefly, DNA oligomers with homology to chromosomal targets and antibiotic resistance cassettes were amplified by PCR from the pKD46 plasmid. The amplified DNA was purified by ethanol precipitation and was then electroporated into the desired strain/species. Cells were allowed to recover in liquid LB at 37 °C for one hour and were then plated on selective media. For clean deletions, a selection and counter-selection method of a tetracycline cassette (selection) with tetracycline-sensitive plates (counter-selection) was used. In these cases, 80-bp ssDNA oligomers were used for the deletion of the tetracycline cassette in the targeted region without PCR amplification. All chromosomal mutations were made with the first and last five intact codons to mitigate potential gene expression polarity artifacts.

In addition to Karlinsey’s method, transduction methods were also used for strain construction in *Salmonella*. For transduction methods, phage P22 was used (*52*).

Plasmids used are summarized in Table S3. All plasmids were constructed using the Gibson Assembly protocols (*53*), and were transformed into *E. coli* and *Salmonella* cells using standard transformation protocols.

### Surface Motility Assays

*Salmonella* surface motility plates were prepared with Luria Broth (LB; 1.0 % Tryptone, 0.50% NaCl, and 0.5 % yeast extract), 0.55 % Difco Agar, and 0.50 % glucose, or other sugar as described in the text. After pouring into Petri dishes, the media was allowed to dry and solidify at room temperature overnight. The filled Petri dishes were then placed into a humidified incubator at 32 °C for 24 hours prior to inoculation. 4 µL of *Salmonella* culture, grown to saturation in liquid LB, was inoculated. The plates were then incubated at 37 °C in approximately 100% humidity.

*E. coli* surface motility plates were prepared similarly. LB, 0.50 % Eiken Agar (Eiken Chem Co. Japan), and 0.50 % glucose were combined. The media was then poured into Petri dishes, and allowed to dry for 8 hours prior to inoculation. 2 µL of *E. coli*, grown to saturation in liquid LB, was inoculated. The inoculated plates were then allowed to dry for 5 minutes in a laminar flow hood, until the inoculant drop had evaporated. The plates were then incubated at 37 °C in 90-100% humidity for 16 hours or as noted otherwise in the text.

### Growth Comparisons

Wells of a Greiner clear-bottom 96-well microplate were filled with 198 µL of LB + 0.50% glucose and were inoculated with 2 µL of saturated culture (1:100), grown in LB media. A lid was placed over the microplate to prevent evaporation, and the microplate was subsequently placed in a BioTek H1 plate reader. The plate reader was set to shake using the orbital shake function at 37 °C and recorded OD_600_ at half-hour intervals for 10 hours. A sigmoidal function was fit to the growth curve data in Python. The doubling time was then extracted from the exponential growth phase of the data.

### Preparation of Bromothymol Blue Solutions and Media Acidification Assay

To examine acidification of the media, we added the pH indicator, bromothymol blue, to the motility assay plates. Bromothymol blue is green at neutral pH, yellow at pH < 6.0, and blue at pH > 7.6. Bromothymol blue stock solution was prepared by adding 100 mg bromothymol blue to 10 mL of 4% NaOH (v/v), to which 10 mL of a 95% (v/v) Ethanol solution was then added. The resulting solution was then further diluted with ddH_2_O to a final volume of 100 mL, yielding a final concentration of 1 mg bromothymol blue/mL. 1.25% (v/v) bromothymol blue solution was added to the media to yield a final concentration of 12.5 µg/mL.

### Microscopy and Cell Tracking

*E. coli* and *Salmonella* strains were grown in liquid cultures to mid-log phase, OD_600_ of 0.4 to 0.6. Cells were pelleted at 1500 g for five minutes and were resuspended in motility buffer (10 mM potassium phosphate, 0.10 mM EDTA, pH 7.5) to a final OD of approximately 0.02. 80 µL of diluted cell culture was pipetted into a tunnel slide (formed by a cover slip fixed on a glass slide with two pieces of double-sided sticky tape). Suspended cells were imaged at 25 fps on a Nikon Eclipse Si phase contrast microscope fitted with a BFS-U3-28S5M-C USB 3.1 Blackfly S, Monochrome Camera. Cell motility videos were then analyzed in ImageJ using Trackmate (*54, 55*).

### Measurements of Acetate and Formate Concentrations within Agar

Plates were sampled by manually extracting agar plugs into a Pasteur pipette, at various positions relative to the margin of an expanding colony. Plugs were combined in a microfuge tube and spun to pellet the agar, then concentrations of acetate or formate in the supernatant were measured using enzyme-based assays (Abcam assay kits ab204719 and ab111748).

## Acknowledgments

We thank Dr. Kelly Hughes and Dr. Fabienne Chevance for sharing strains and for extensive discussions, Dr. Souvik Bhattacharyya and Dr. Siddarth Srinivasan for advice, Eliza Kunzler for assistance with measurements of swarming and swashing, and the members of Wadhwa and Blair labs for fruitful discussions. This work was partially supported by the National Institute of General Medical Sciences of the National Institutes of Health under Award Number R00GM134124 (to N.W.).

## Data and materials availability

All data are available in the main text or the supplementary materials.

