## Supplemental Material for "Swashing motility: A novel propulsion-independent mechanism for surface migration in *Salmonella* and *E. coli*"

**This PDF file includes:**

Figs. S1 to S13
Tables S1 to S3
Reference (56)

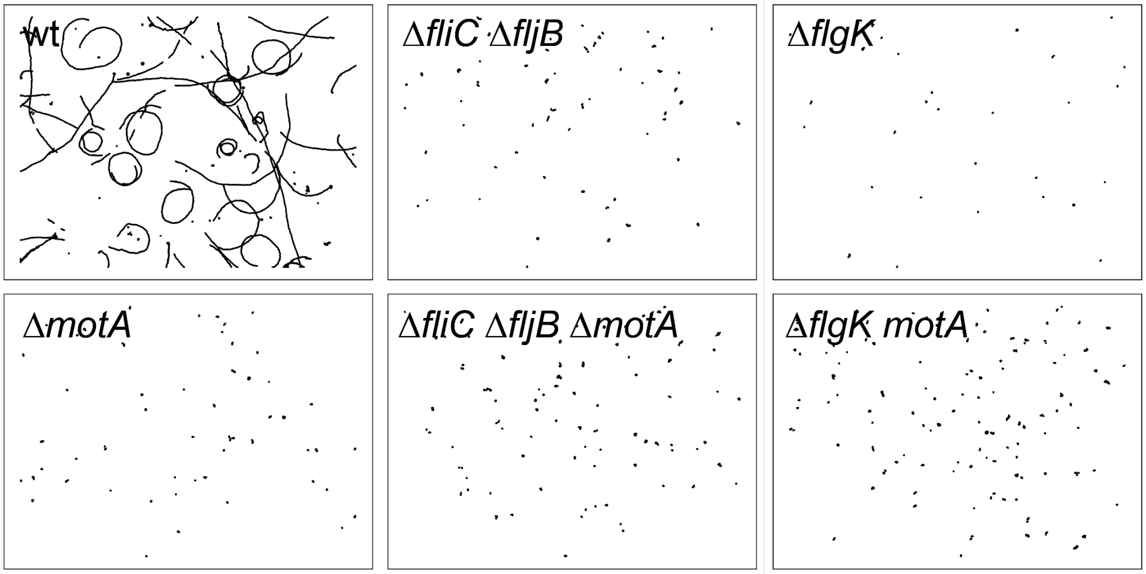


**Fig. S1.**

***Salmonella* microscopy and cell tracking.** WT cells swim using run and tumble motion. In contrast, cells lacking either the filament (Δ*fliC* Δ*fljB;* Δ*flgK;* Δ*flgL*) or the stator proteins (Δ*motA;* Δ*motB*) do not swim. The swimming patterns seen for wt cells are consistent with clockwise circular motion reported for cells interacting with the boundary, in this case, the glass coverslip (*56*).

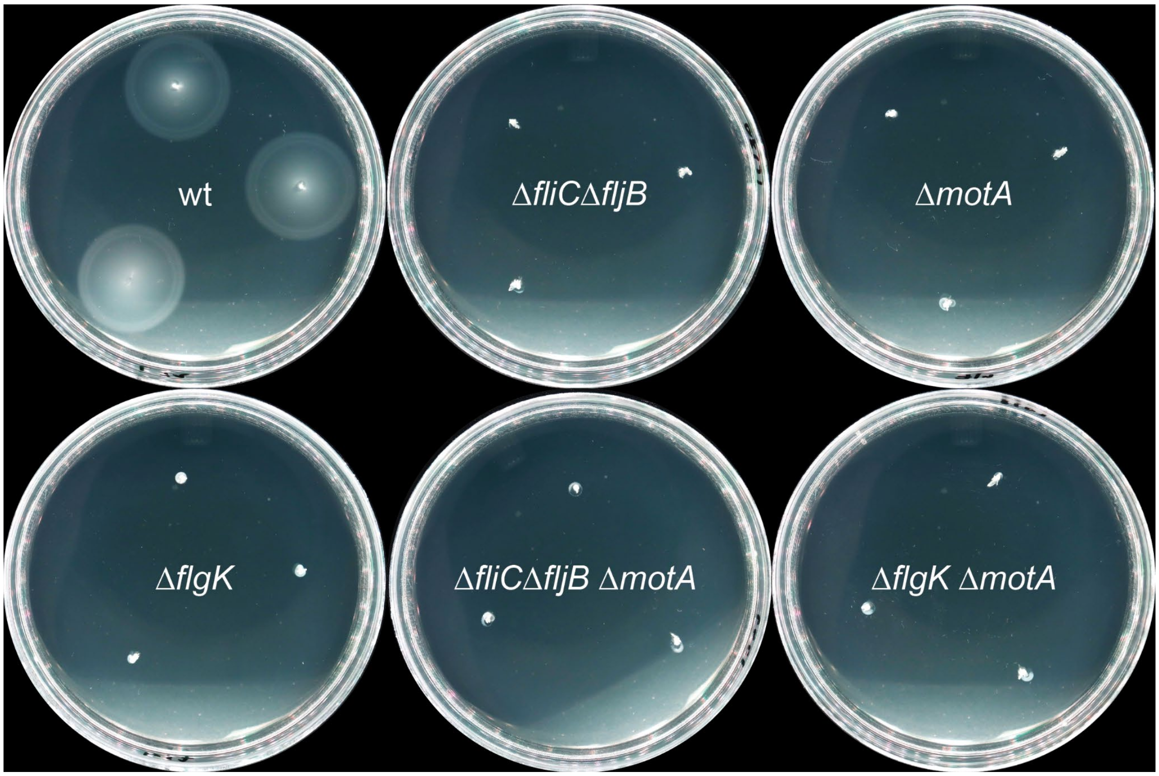


**Fig. S2.**

***Salmonella* motility assays in 0.30% agar swim plates.** WT cells migrate outwards, showing swimming behavior. In contrast, cells lacking either the filament (Δ*fliC* Δ*fljB;* Δ*flgK;* Δ*flgL*) or the stator proteins (Δ*motA*) are unable to migrate outwards. Each plate shows three biological replicates.

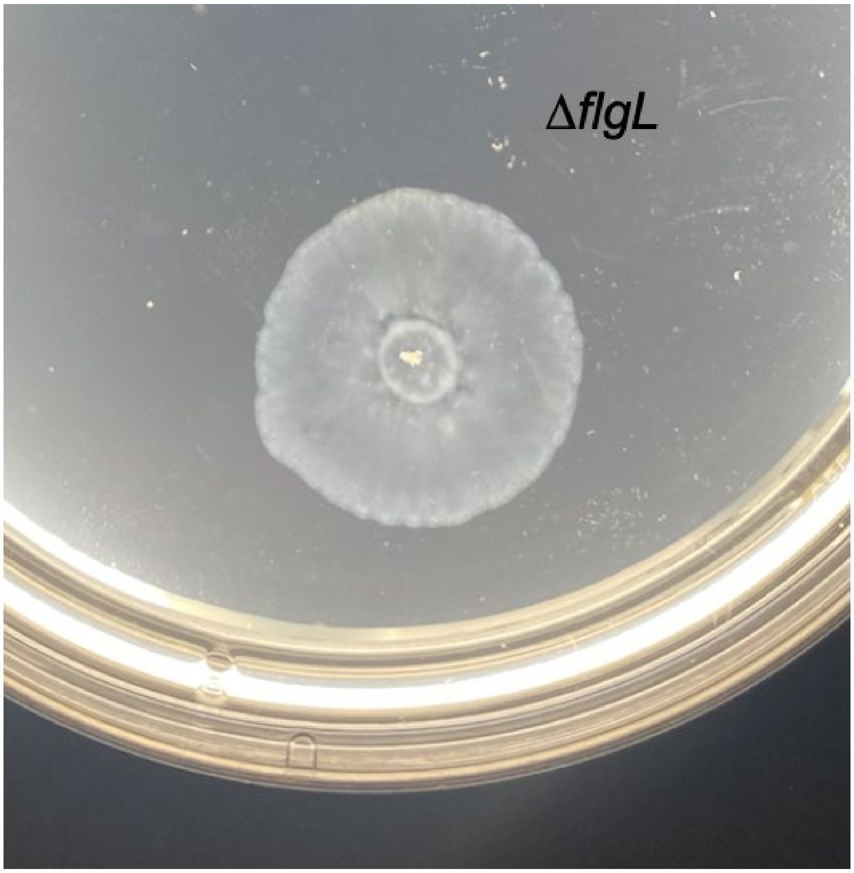


**Fig. S3.**
Migration of *Salmonella* Δ*flgL* cells on a swarm plate, imaged at 8 hours.

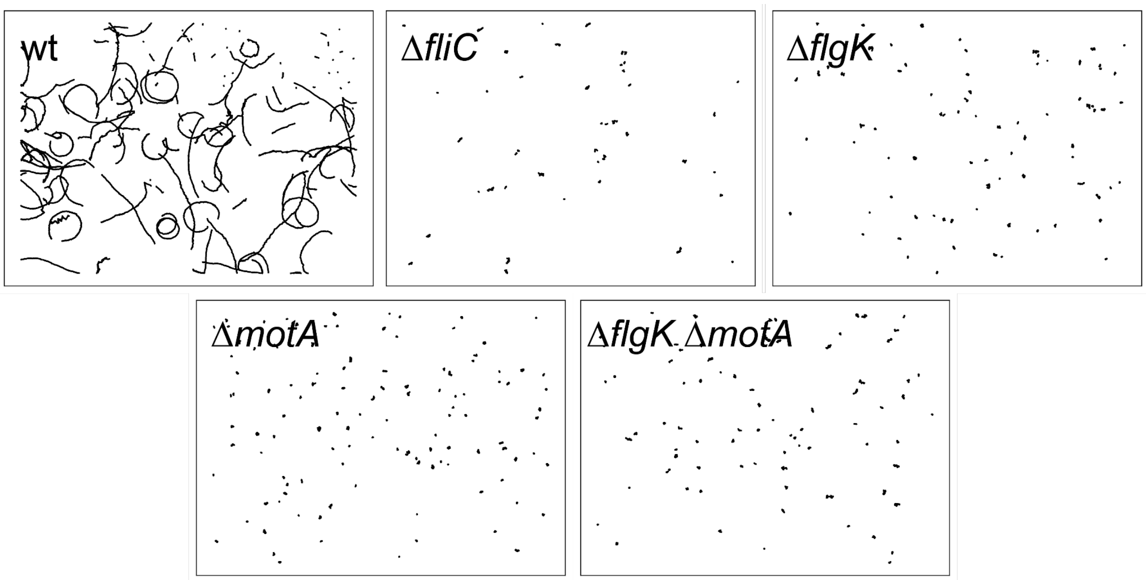


**Fig. S4**.

***E. coli* microscopy and cell tracking.** WT cells swim using run and tumble motion. In contrast, cells lacking either the filament (Δ*fliC;* Δ*flgK*) or the stator proteins (Δ*motA;* Δ*motB*) or both (Δ*motA*Δ*flgK*) are unable to swim. The swimming patterns seen for wt cells are consistent with clockwise circular motion reported for cells interacting with the boundary, in this case, the glass coverslip (*56*).

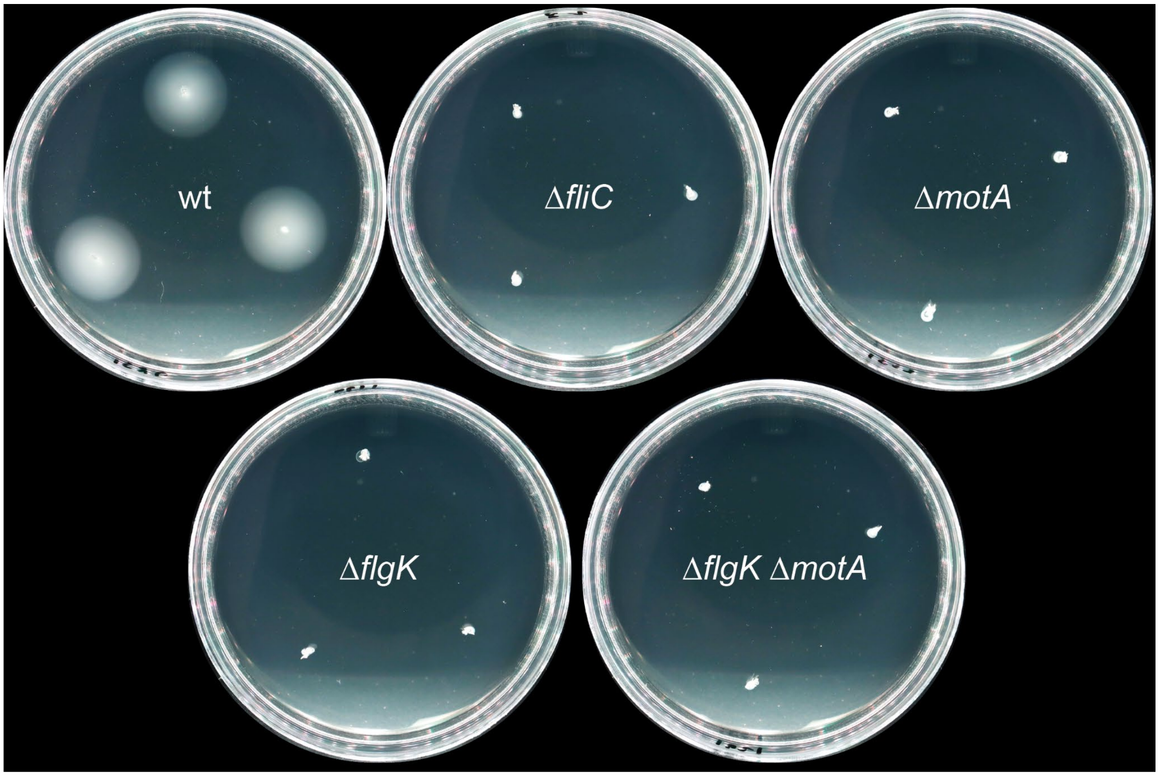


**Fig. S5**.

***E. coli* motility assays in 0.30% agar swim plates.** WT cells migrate outwards, showing swimming behavior. In contrast, cells lacking either the filament (Δ*fliC;* Δ*flgK*), the stator proteins (Δ*motA*), or both (Δ*motA*Δ*flgK*) are unable to swim. Each plate shows three biological replicates.

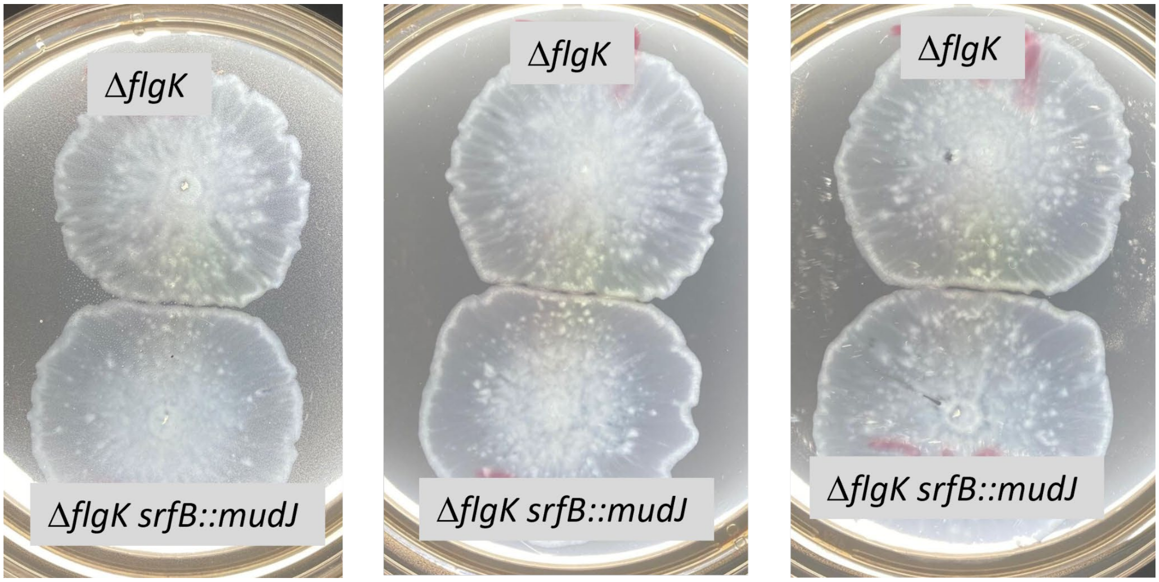


**Fig. S6.**

Swashing of the *Salmonella ΔflgK* mutant is not prevented by loss of SrfB. The image shows three biological replicates; plates were photographed after 10.5 h at 37° C.

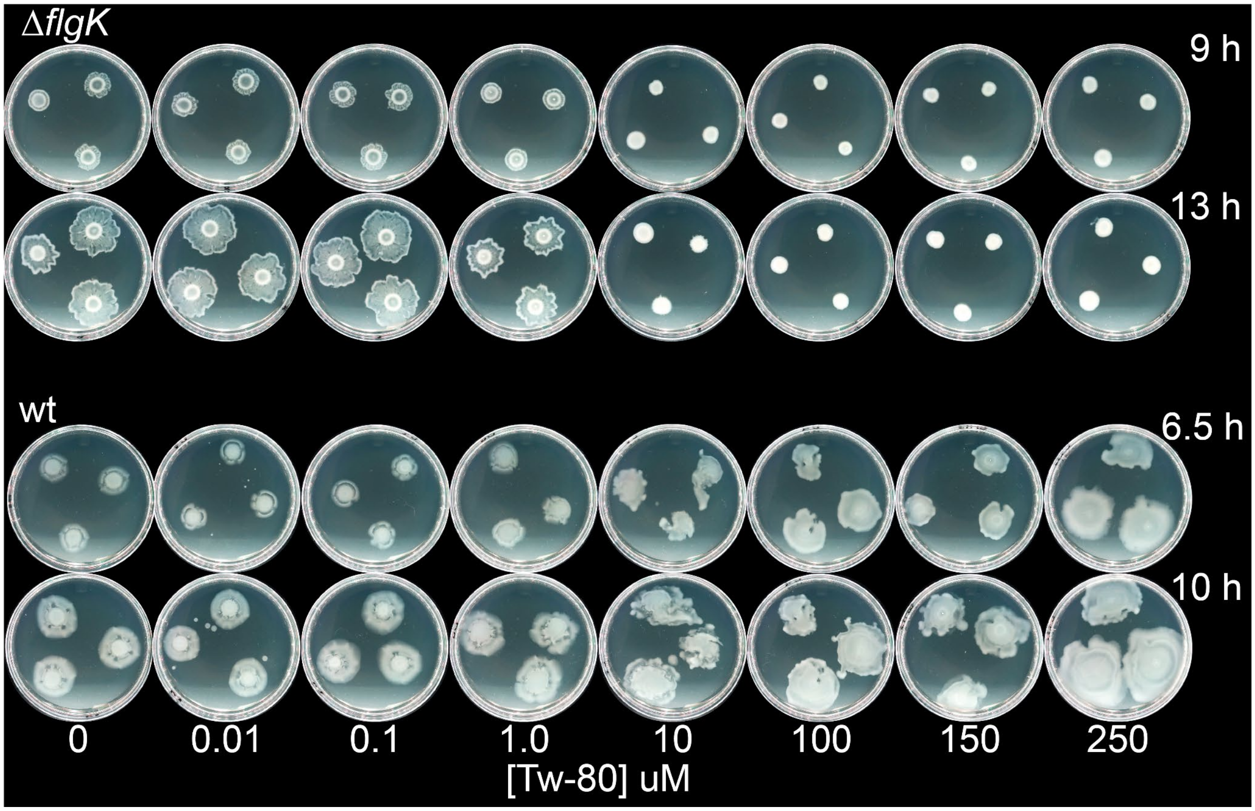


**Fig. S7.**

Inhibition of *E. coli* swashing, and enhancement of swarming, by surfactant, Tween-80.

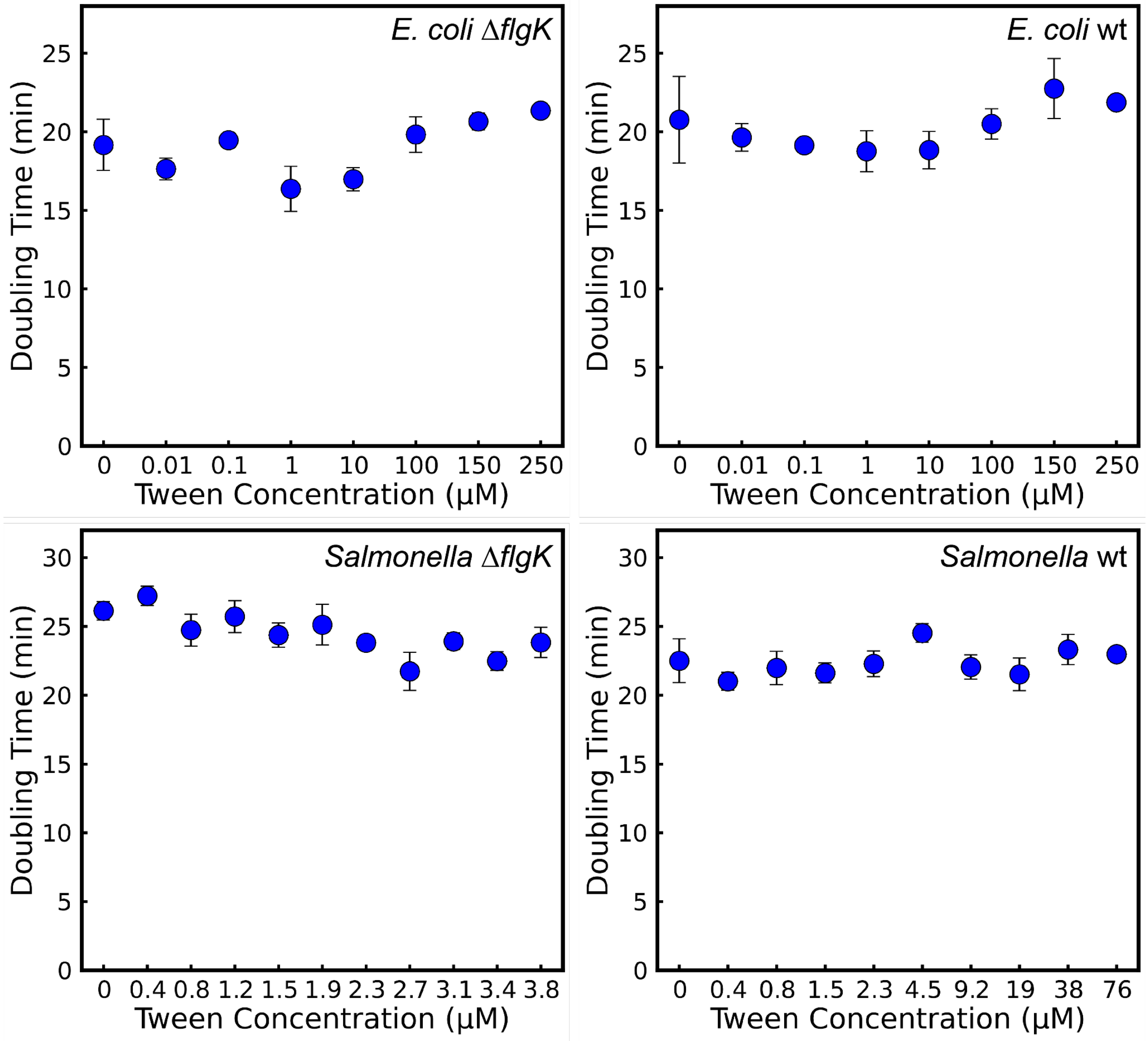


**Fig. S8.**

Addition of Tween-80 does not affect *Salmonella* or *E. coli* growth.

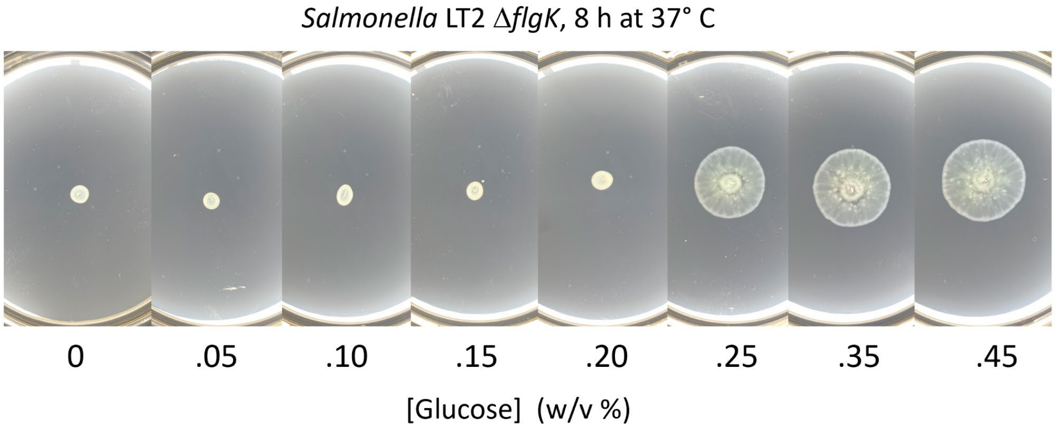


**Fig. S9.**

Surface migration of *Salmonella* Δ*flgK* strain ceased at glucose concentrations below 0.25%.

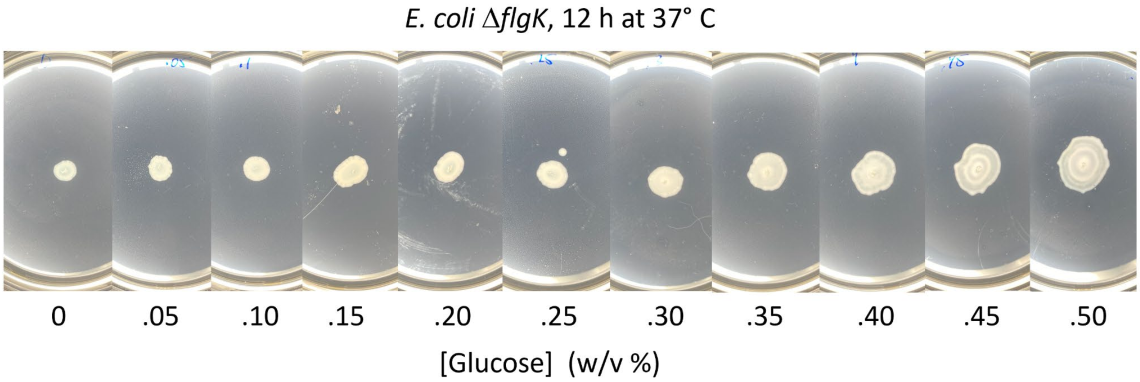


**Fig. S10.**
Surface migration of *E. coli* Δ*flgK* strain ceased at glucose concentrations below 0.3%.

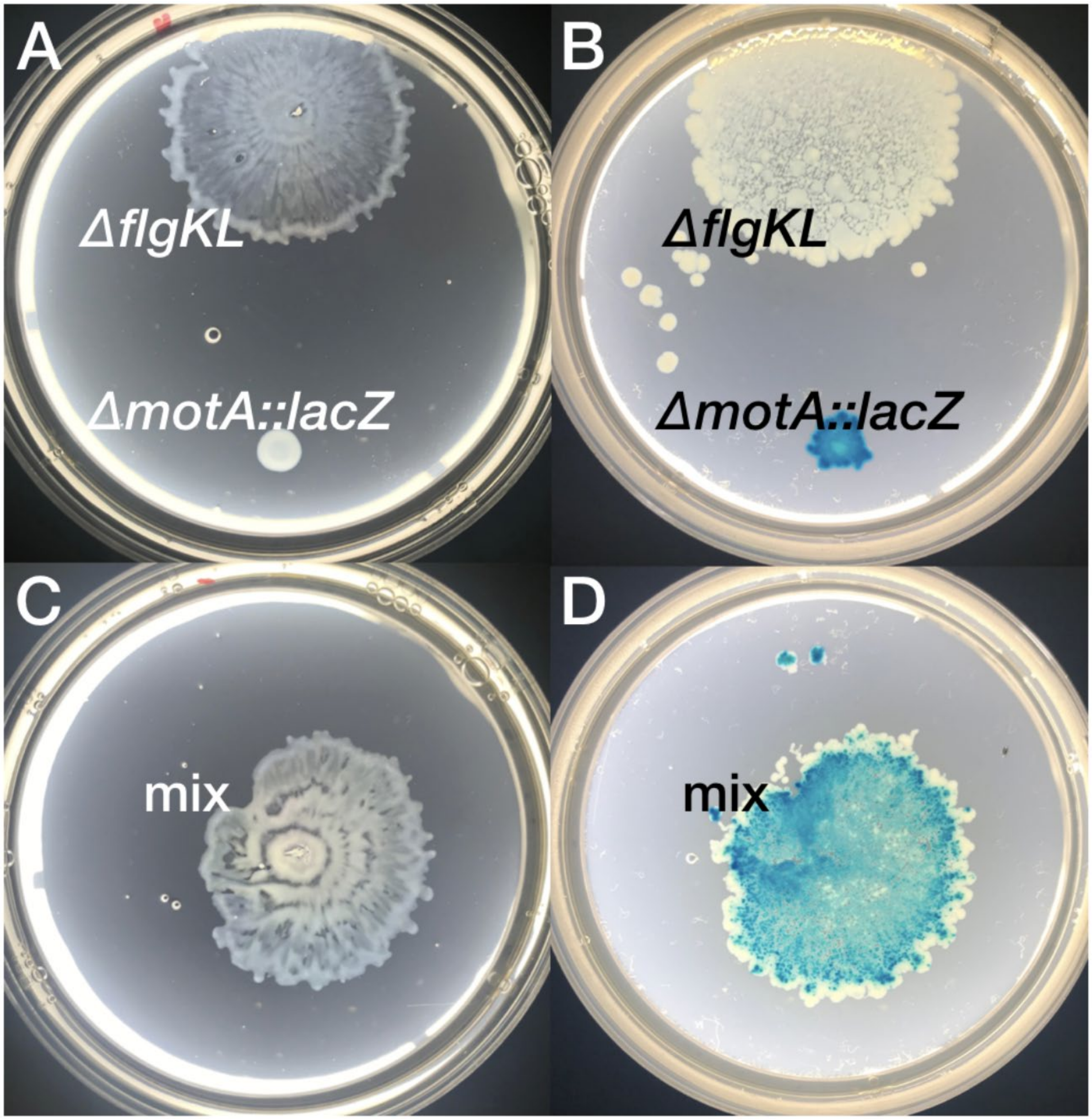


**Fig. S11.**

*Salmonella* ∆*motA* shows diminished migration relative to ∆*flgKL* (*A*), whereas a 1:1 mixture of ∆*motA* with ∆*flgKL* demonstrates uninhibited expansion (*C*). On X-galactose plates, a lacZ knock-in of ∆*motA* appears blue (*B*) and migrates to the edge of the swashing mixture alongside ∆*flgKL* (*D*).

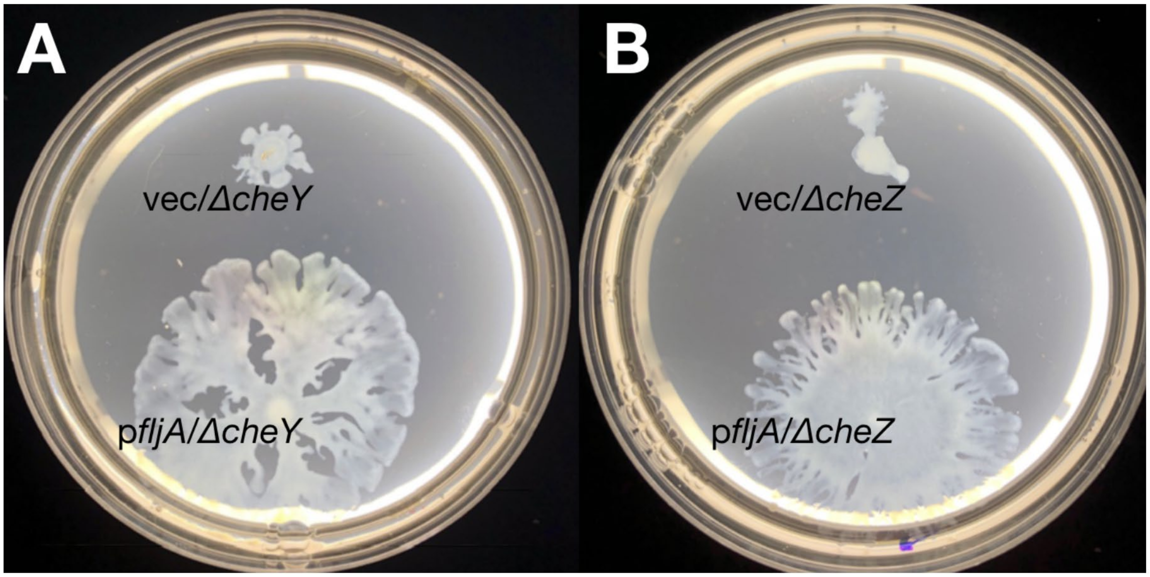


**Fig. S12.**

Surface motility of chemotactic knockout strains, ∆*cheY (A)* and ∆*cheZ (B)*, is recuperated through further inhibition of flagellin protein translation via expression of *fljA*.

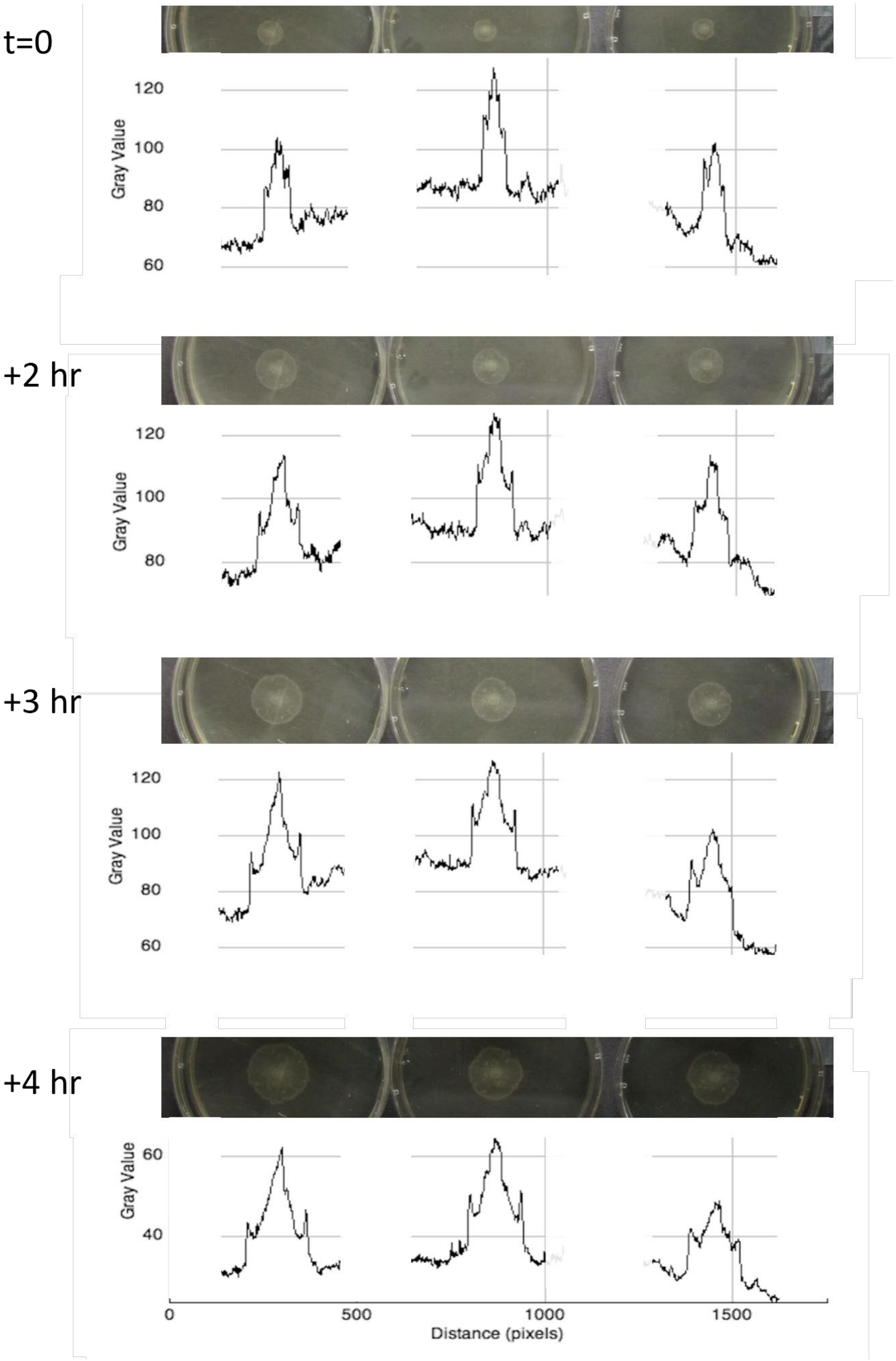


**Fig. S13.**

Densitometry measurements at the margin of swashing ΔflgK colonies. Each row represents 3 biological replicates, and four different time points are shown.

| Strain Name | Background | Genotype | Source |
|---|---|---|---|
| JP1 | *S. enterica* LT2 | w.t. | K. Hughes |
| JP719 | *S. enterica* LT2 | Δ*flgK2408* | K. Hughes |
| JP720 | *S. enterica* LT2 | Δ*flgL2403* | K. Hughes |
| JP799 | *S. enterica* LT2 | Δ*fliC* Δ*fljB* | K. Hughes |
| JP1049 | *S. enterica* LT2 | Δ*motA::tetRA* Δ*fliC* Δ*fljB* | This study |
| JP1057 | *S. enterica* LT2 | Δ*motA::tetRA* | This study |
| JP1058 | *S. enterica* LT2 | Δ*motA::tetRA* Δ*flgK2408* | This study |
| JP1117 | *S. enterica* LT2 | Δ*flgKL* | K. Hughes |
| TH29200 | *S. enterica* LT2 | Δ*flgK ackA::mudJ* | This study |
| DBS710 | *S. enterica* LT2 | Δ*flgK srfB::mudJ* | This study |
| JP1533 | *S. enterica* LT2 | Δ*motA::lacZ* | This study |

**Table S1.**
*Salmonella* strains used in this study.

| Strain Name | Background | Genotype | Source |
|---|---|---|---|
| JP1586 | *E. coli K12 MG1655 F- lambda- mot+* | w.t. | K. Hughes |
| JP1593 | *E. coli K12 MG1655 F- lambda- mot+* | Δ*motA::tetRA* | This study |
| JP1594 | *E. coli K12 MG1655 F- lambda- mot+* | Δ*flgK::tetRA* | This study |
| JP1621 | *E. coli K12 MG1655 F- lambda- mot+* | Δ*flgK* Δ*motA::tetRA* | This study |
| DFB28 | *Escherichia coli* RP437 | Δ*fliC* | This study |

**Table S2.**

*E. coli* strains used in this study.

| Plasmid Name (resistance) | Clone | Source |
|---|---|---|
| pKD46 (ApTs) | Lambda red proteins | D. Blair |
| pMS421 (Sp) | vector | K. Hughes |
| pJP6 (Sp) | pMS421::*fljA* | This study |

**Table S3**.
Plasmids used in this study.
